# Increased Expression and Altered Functional Activities of Immune Receptors TREM1, PD-L1, and Others on Hematopoietic Progenitor Cells in a Mouse Model of Rheumatoid Arthritis

**DOI:** 10.64898/2026.06.11.731762

**Authors:** Johanna Toth, Roselyn R. Jiang, Lin Tze Tung, Dania Shaban, Mathieu Mancini, Bianca Pozzebon, Joo Eun (June) Kim, Mitra Yousefi, Danielle Malo, Silvia M. Vidal, Ines Colmegna, David Langlais, Anastasia Nijnik

## Abstract

Hematopoietic stem and progenitor cells (HSPCs) sustain the production of hundreds of billions of new cells per day to maintain our blood and immune system. In this process, HSPCs regulate the hematopoietic output by sensing and integrating diverse physiological cues. Thus, HSPCs express many receptors traditionally studied for their functions in the immune system, and this allows HSPCs to directly detect microbial compounds, endogenous danger signals, cytokines, and other inflammatory mediators. However, how the expression levels of such receptors on HSPCs change under chronic inflammation and how such changes alter HSPC functions and immune cell production remains unexplored. Working in a murine model of rheumatoid arthritis, we demonstrate the induction of microbial sensors TLR2 and CD14, orphan inflammatory receptor TREM1, and checkpoint receptor PD-L1 on HSPCs and particularly the myeloid progenitor cells in the arthritis-afflicted mice. Furthermore, we demonstrate that the stimulation of HSPCs through these receptors in culture can significantly alter the dynamics of cell expansion and differentiation, with distinct responses from HSPCs of arthritis-afflicted versus healthy control mice. We hypothesize that the induction and stimulation of HSPCs through these immune receptors under chronic inflammation may impact the output and functional properties of their immune cell progeny, positing HSPCs as central players in the pathogenic inflammatory responses of rheumatoid arthritis and potentially other chronic inflammatory diseases.

**HIGHLIGHTS:**

- Hematopoietic progenitor cells in murine models of rheumatoid arthritis show an upregulation of immune receptors TREM1, PD-L1, TLR2, and CD14.
- Stimulation of murine hematopoietic stem and progenitor cells through these receptors in culture alters the dynamics of their expansion and differentiation.
- In such cultures, hematopoietic stem and progenitor cells from mice afflicted with rheumatoid arthritis show altered responses to stimulation as compared to healthy controls.

## Introduction

Hematopoietic stem and progenitor cells (HSPCs) sustain the production of hundreds of billions of new cells per day to maintain our blood and immune system. In this process, HSPCs regulate the hematopoietic output by sensing and integrating diverse physiological cues [1-3]. In response to acute inflammation, HSPCs increase their proliferation and myeloid-biased differentiation, ensuring the rapid production of myeloid leukocytes for defense against infections [4]. In contrast, chronic inflammation can drive progressive HSPC dysfunction, associated with oxidative stress, transcriptional and metabolic reprogramming, accumulation of genetic lesions, and telomere attrition [5-8]. A prominent example of this is rheumatoid arthritis (RA) – chronic systemic autoimmune and inflammatory disease that affects up to 1% of the adult population [9, 10]. Multiple recent studies characterize HSPC dysfunction in RA and its impact on the immune cell production, in both human patients [7, 8, 11] and murine models [12-15].

HSPCs express many pattern recognition receptors (PRRs) as well as other immune receptors that allow the direct detection of microbial compounds, endogenous danger signals, cytokines, and other inflammatory mediators produced by our immune system [1-3, 5]. Thus, HSPC responses to many classical microbial and endogenous inflammatory signals have been characterized ex vivo and following an acute challenge in murine models [1, 16]. However, the expression and function of many other immune receptors on HSPCs remain fully unexplored. One such example is the orphan receptor TREM1, strongly implicated in the immunopathology of many systemic inflammatory diseases [17-21]. Checkpoint receptor and drug-target PD-L1 was shown to be expressed by HSPCs [22, 23], however its functional role in HSPCs or hematopoiesis has not been investigated [24]. Furthermore, how the steady state levels of immune receptor expression on HSPCs change under chronic inflammation and how such changes may alter HSPC functions and immune cell production remains fully unexplored.

Here, we assess the cell-intrinsic role of select immune receptors, including TLR2, TREM1, and PD-L1 on HSPCs in a mouse model of RA, as the prototypic chronic inflammatory disease. We test how systemic inflammation in murine RA affects the expression of these immune receptors on HSPCs, and how the altered receptor expression impacts HSPC expansion, differentiation, and responses to inflammatory signals.

## Materials and Methods

### Mice, Flow Cytometry, and RNA-seq Analyses

SKG mouse model of rheumatoid arthritis is widely used [13, 25], including in our recent work [26]. Flow cytometry and RNA-seq analyses methods were previously described [27, 28], with full information and antibodies provided in Supplemental.

### HSPC Cultures

HSPCs were isolated from murine bone marrow using EasySep CD117 Positive Selection Kit (Stem Cell Technologies, 18757). HSPCs were cultured in media made with equal parts IMDM and DMEM (Life Technologies, 12440-053 & 11995-065) with 10% HyClone Bovine Growth serum (Fisher, SH30541.03), 2mM L-glutamine (Life Technologies, 609-065-EL), 0.2μM β-mercaptoethanol, 5ng/mL IL-3 (200-03), 10ng/ml IL-6 (200-06), 100ng/mL SCF (300-07), 100ng/mL TPO (300-18), and 100ng/mL FLT-3L (300-19), all from PeproTech, ThermoFisher Scientific. HSPCs were stimulated with (1→3)-β-D-glucan (*Alcaligenes faecalis*, Millipore-Sigma, 5μg/mL), lipopolysaccharide (LPS, *E. coli* O111:B4, Millipore-Sigma, 10ng/mL), or by pre-coating wells with 10μg/mL anti-TREM-1 agonist antibody (Biotechne, MAB1187), anti-PD-L1 blocking antibody (eBioscience, MIH5, 16-5982-81), or rat IgG2a isotype control (eBioscience, eBR2a, 16-4321-81).

### Human subjects

Patient recruitment criteria were: age >18, rheumatoid arthritis classification according to 2010 American College of Rheumatology & European League Against Rheumatism criteria, seropositive for rheumatoid factor and/or ACPA, with no coexisting systemic inflammatory disorders, infectious diseases, or cancer. Healthy controls matched by age and sex were also recruited. Participants provided written informed consent. Demographic characteristics of patients are provided in Table S4.

### Ethics

Mouse experiments were in accordance with guidelines of Canadian Council on Animal Care and protocols MCGL-7932/6029 approved by McGill Animal Care Committee. Studies with human subjects were approved by the McGill University Health Center Ethics Review Board (ERB: 09-239 GEN), and written consent obtained from all participants.

## Results and Discussion

### Induction of TREM1, TLR2, and CD14 on myeloid progenitors in RA

To assess how chronic systemic inflammatory disease affects immune receptor expression on HSPCs, we analyzed HSPCs from murine SKG model of RA [13, 25, 26]. Early myeloid progenitor cells from RA-afflicted mice showed significantly increased levels of TREM1, TLR2, and CD14, but reduced levels of CLEC7A/Dectin-1, relative to HSPCs of healthy control SKG mice (Figure 1A-B), with no changes observed for earlier stem cells and multipotent progenitor cells (HSC-MPPs, not shown). Analyses of CD34^+^ HSPCs in the blood of RA patients similarly indicated upregulation of TREM1, TLR2, and CD14 (Figure S1).

**Figure 1.**
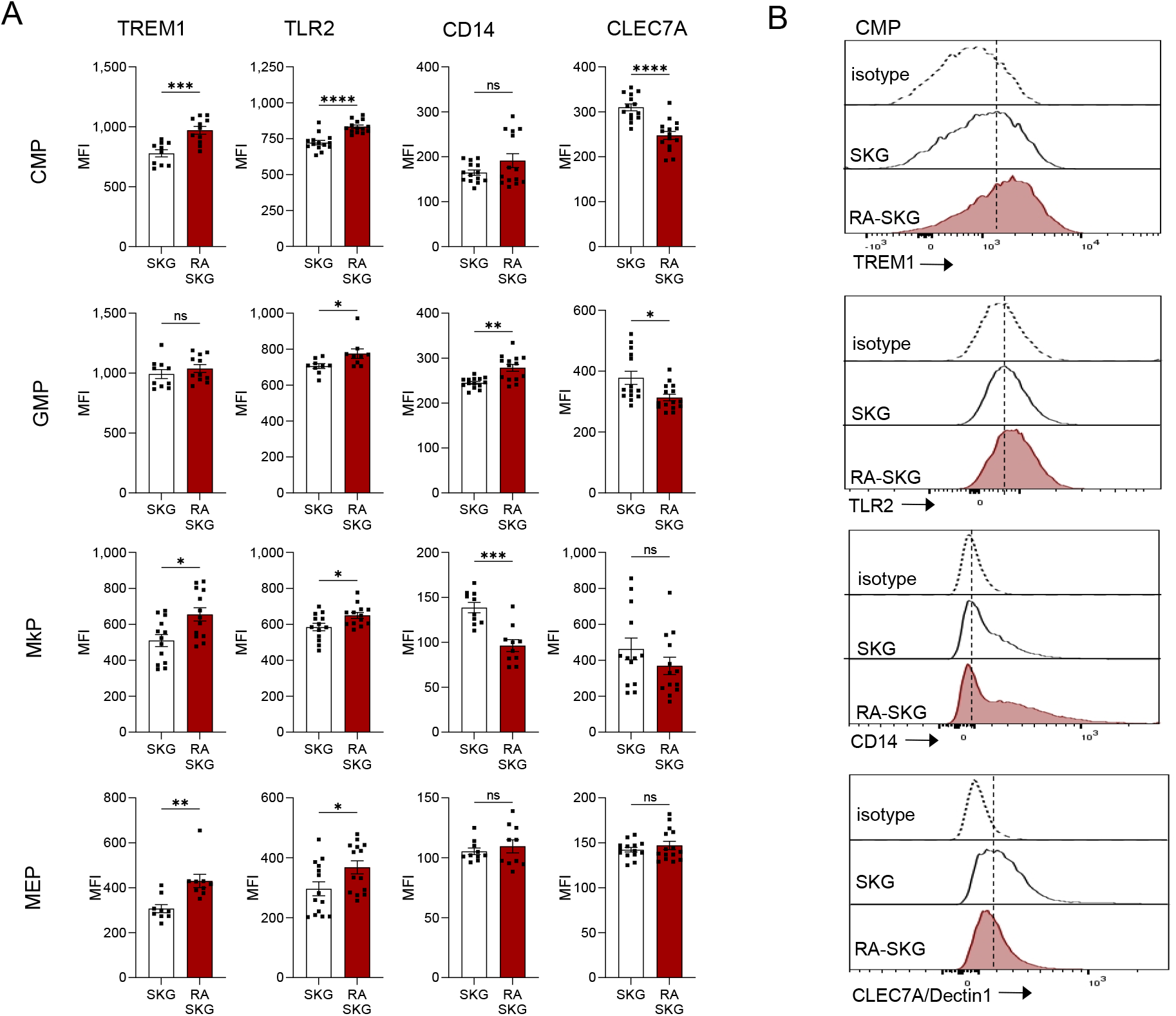
Increased levels of immune receptors TREM1, TLR2, and CD14 on hematopoietic progenitor cells in a murine model of rheumatoid arthritis. **(A)** Cell-surface levels of TREM1, TLR2, CD14, and CLEC7A (Dectin-1) on HSPCs of RA-afflicted and healthy control SKG strain mice, analyzed by flow cytometry of bone marrow. The cells presented are common myeloid progenitors (CMPs), granulocyte monocyte progenitors (GMPs), megakaryocyte progenitors (MkPs), and megakaryocyte erythroid progenitors (MEPs), gated as Lin^−^cKit^+^Sca1^−^ followed by CD34^+^CD16/32^−^ for CMPs, CD34^+^CD16/32^+^ for GMPs, CD34^−^CD16/32^−^ for MEPs, and CD41^+^CD150^+^ for MkPs. **(B)** Representative flow cytometry histograms of immune receptor expression on CMPs; isotype control antibodies were used to confirm positive staining. Data are from 5-15 mice per group, consolidated from 2-3 independent experiments. Bars represent means ± SEM; statistical analyses used Mann-Whitney test; ^*^ *p*<0.05, ^**^ *p*<0.01, ^***^ *p*<0.001, ^****^ *p*<0.0001, ns – not significant; MFI – mean fluorescence intensity.

### Altered responses of RA-HSPCs to microbial ligands

To test how the altered expression of immune receptors on RA-HSPCs affects cellular functions, HSPC culture model was adopted, as in vivo systems do not easily allow selective HSPC stimulation or selective disruption of receptor expression on HSPCs without affecting their downstream immune cell progeny. HSPCs from healthy control and RA-afflicted mice were cultured for 18 hours with TLR2/Dectin-1 ligand β-glucan and with TLR4/CD14-ligand LPS. In cultures of healthy control HSPCs, both microbial ligands strongly suppressed the maintenance and expansion of all HSPC subsets, from stem cells and earliest multipotent progenitors HSC-MPP1, to myeloid-biased multipotent progenitors MPP2-MPP3, and the myeloid-committed progenitors CMP-GMP (Figure 2A).

**Figure 2.**
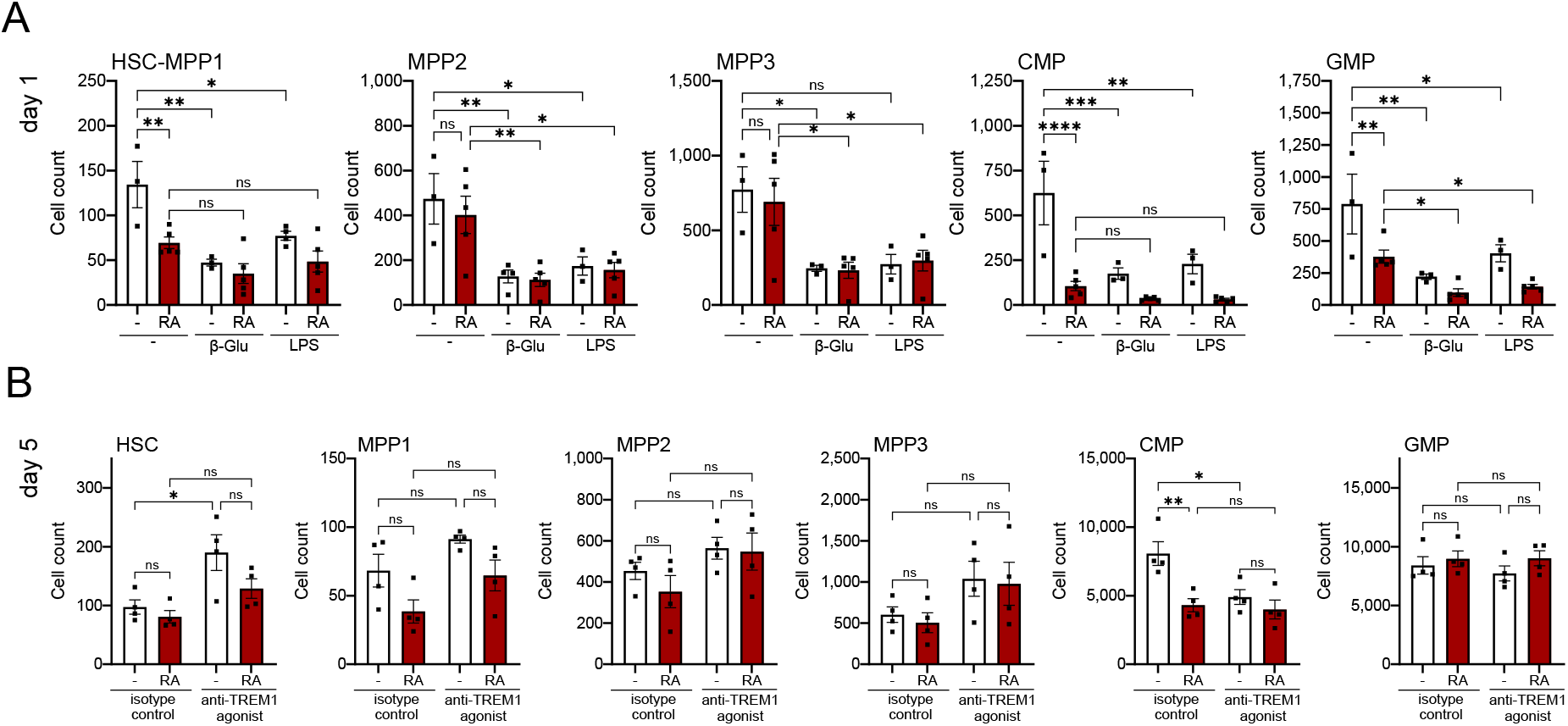
Altered responses of RA-HSPCs to stimulation with microbial ligands and anti-TREM1 agonist antibody. The cells analyzed are HSCs – hematopoietic stem cells, MPPs – multipotent progenitor cells, CMPs – common myeloid progenitors, and GMPs – granulocyte-monocyte progenitors, gated as shown in Figure S3. **(A)** Analyses of HSPCs from RA-afflicted versus healthy control SKG strain mice, cultured for 18 hours with the TLR2/Dectin-1 ligand β-glucan at 5μg/mL, or with the TLR4/CD14 ligand LPS at 10ng/mL, with the HSPC subsets quantified by flow cytometry. **(B)** Analyses of HSPCs from RA-afflicted versus healthy control SKG strain mice, cultured for 5 days in wells pre-coated with an anti-TREM1 agonist antibody or an isotype control antibody, with the HSPC subsets quantified by flow cytometry. Bars represent means ± SEM; datapoints represent individual mice at n=4 per group, with data representative of 1-2 independent experiments; statistical analyses used ANOVA with Sidak’s post-hoc test; ^*^ *p*<0.05, ^**^ *p*<0.01, ^***^ *p*<0.001, ^****^ *p*<0.0001, ns – not significant.

Relative to untreated healthy control HSPCs, RA-HSPC cultures showed impaired expansion of HSC-MPP1 and CMP-GMP cells (Figure 2A), which contrasts with the in vivo induction of emergency myelopoiesis in RA [12-15]. Microbial stimulation of RA-HSPCs resulted in further downregulation in the expansion of MPP2, MPP3, and GMP cells, with no significant effects on other subsets. Overall, given the reduced expansion of RA-HSPCs in control untreated cultures, the repressive effects of microbial stimulation on these cells appeared diminished, despite increased receptor expression. This may indicate RA-HSPC dysfunction due to their chronic inflammatory overstimulation in situ.

### Distinct responses to TREM1 stimulation in cultures of healthy control and RA-HSPCs

To test if HSPCs can directly respond to TREM1-stimulation and how such responses are altered in RA, HSPCs were plated into wells pre-coated with anti-TREM1 agonist antibody and the cultures analyzed at day 5. TREM1-stimulation of HSPCs from healthy control mice promoted the expansion of HSCs but inhibited the expansion of CMPs (Figure 2B). In contrast to healthy control HSPCs, RA-HSPCs showed no significant responses to TREM1 stimulation (Figure 2B). This is unexpected given the increased TREM1 expression on RA-HSPCs, but may reflect previous chronic stimulation of the cells in situ that acts to desensitize them to such secondary stimulation.

### Immune receptor induction on RA-HSPCs involves increased gene expression

To establish whether induction of immune receptors on RA-HSPCs involves increased gene expression, we reanalyzed RNA-seq data of Regan-Komito et. al. from SKG mice with chronic systemic inflammation (GSE126218) [13]. This revealed increased *Trem1* transcript in MPP3-4 and *Cd14* transcript in HSCs and MPP3-4 cells. While *Tlr2* transcript was not significantly increased, there was increased expression of other TLR-encoding genes in HSC-MPPs (Figure S2). Furthermore, systemic inflammation resulted in increased *Cd274* transcript levels in HSCs and MPPs, encoding checkpoint receptor PD-L1 (Figure S2).

### Increased PD-L1 levels and responses to PD-L1 blockade in cultures of RA-HSPCs

We validate increased expression of PD-L1 on HSPCs from RA-afflicted mice with flow cytometry, affecting all subsets from HSCs to myeloid progenitors (Figure 3A-B). In contrast, there was no significant PD-L1 induction on lymphoid-primed MPP4 or megakaryocyte-erythroid MEP progenitors (not shown).

**Figure 3.**
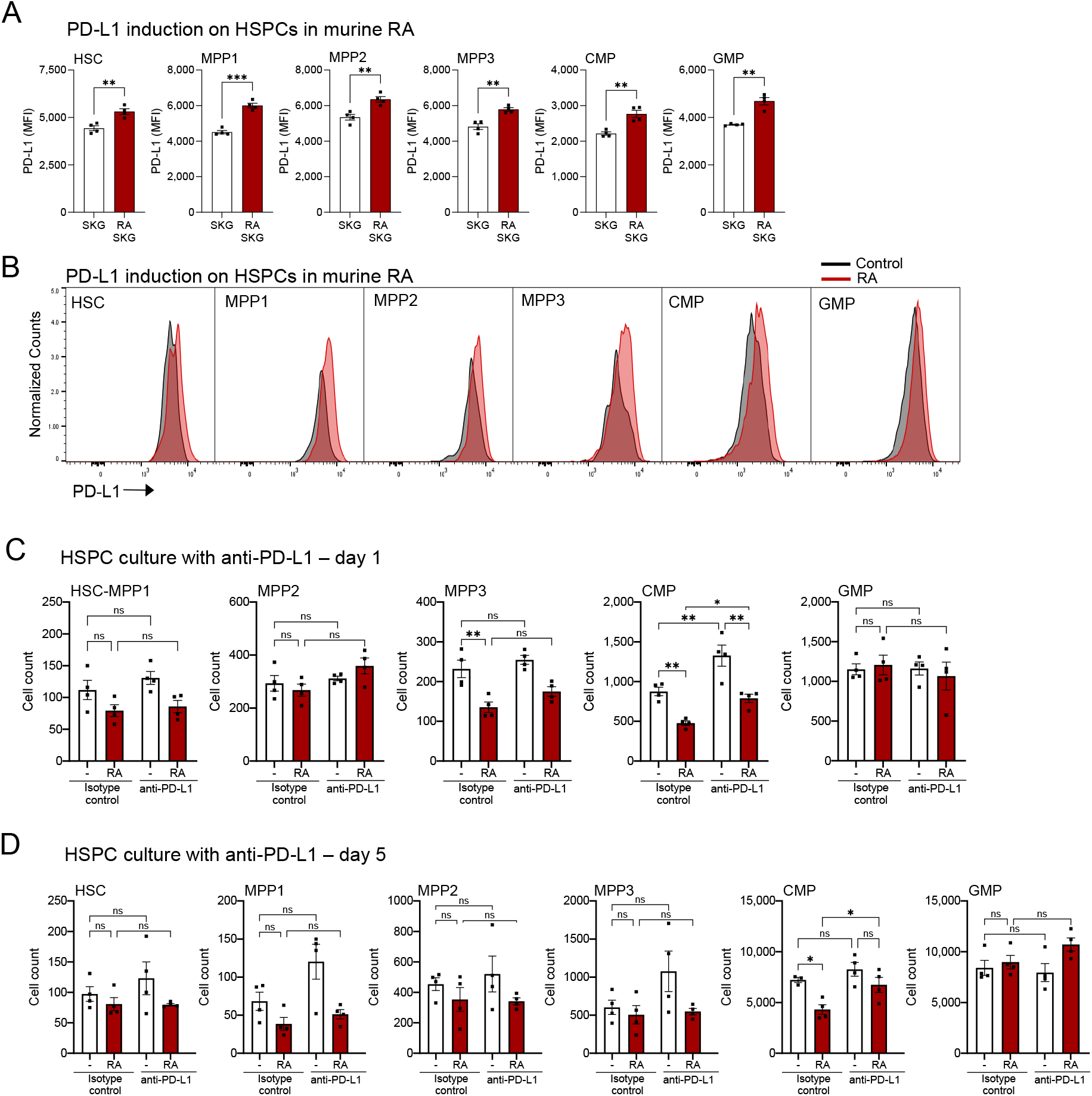
Increased PD-L1 levels on RA-HSPCs and altered responses to PD-L1 blockade in RA-HSPC cultures. The cells analyzed are HSCs – hematopoietic stem cells, MPPs – multipotent progenitor cells, CMPs – common myeloid progenitors, and GMPs – granulocyte-monocyte progenitors, gated as shown in Figure S3. **(A-B)** Increased levels of PD-L1 on HSPC subsets of arthritis-afflicted relative to healthy control SKG strain mice, measured via flow cytometry of bone marrow. **(C-D)** Analyses of HSPCs from RA-afflicted versus healthy control SKG strain mice, cultured for 1 or 5 days in wells pre-coated with an anti-PD-L1 blocking antibody or an isotype control antibody, with the HSPC subsets quantified by flow cytometry. Bars represent means ± SEM; datapoints represent individual mice, with n=4 per group; statistical analyses used (A) unpaired *t*-test with Welch’s correction, or (C-D) ANOVA with Sidak’s post-hoc test; ^*^ *p*<0.05, ^**^ *p*<0.01, ^***^ *p*<0.001, ns – not significant; MFI – mean fluorescence intensity.

PD-L1 blockade in HSPC cultures promoted the expansion of CMPs, with only minor effects on other cell subsets, and this was observed for HSPCs from both healthy and RA-afflicted mice (Figure 3C-D). As already discussed, untreated cultures of RA-HSPCs consistently produced fewer CMPs than cultures of healthy control HSPCs (Figure 2-3). Importantly, PD-L1 blockade diminished this effect by day 1 and fully abolished it by day 5, allowing the normal expansion of CMPs in RA-HSPC cultures (Figure 3C-D). This indicates that at least in this ex vivo system PD-L1 blockade can antagonize and reverse the effects of chronic systemic inflammation on HSPCs.

### Discussion

We demonstrate altered expression of immune receptors on HSPCs, and particularly myeloid progenitors, in arthritis-afflicted mice, and apply cell culture systems to analyze the distinct responses of RA-HSPCs to microbial, inflammatory, and danger signals.

We establish that HSPCs can directly respond to stimulation of TREM1 – orphan receptor strongly implicated in immunopathology of chronic inflammatory diseases, including RA [17-21]. This raises important questions regarding TREM1 contribution to the induction of emergency myelopoiesis, and whether its cell-intrinsic functions on HSPCs may affect the course of disease progression. We further show that PD-L1 blockade promotes myeloid progenitor cell expansion in HSPC cultures, and over time reverses some features of RA-HSPC dysfunction.

Inherent limitation of the study is the use of HSPC cultures, due to the challenges of selective HSPC modulation in situ without affecting their niche and immune cell progeny. Thus, future studies should explore the cell-types and ligands that stimulate HSPCs via TREM1 and PD-L1 in situ, signaling pathways induced by these receptors in HSPCs [21, 29, 30], and their possible roles in HSPC functions as antigen presenting cells in the context of other diseases, including hematologic malignancies and their immunotherapy [24].

Overall, this study posits HSPCs as central players in inflammatory responses and novel entry-points to understand and target immune dysregulation.

## Acknowledgments

Work was funded by Canadian Institutes of Health Research Project Grant (CIHR PJT-173417, to AN, DL, IC), Team Grant from McGill Interdisciplinary Initiative in Infection & Immunity (MI4) through generous gift from the Elizabeth Duthie-Saunders Estate (to SMV, DL, AN, DM, IC), and Internal Award from McGill Regenerative Medicine Network (to IC, AN). AN was a Canada Research Chair (CRC) Tier II in Hematopoiesis; SMV is a CRC Tier I in Host Responses to Virus Infections; DL is a Research Scholar of the Fonds de Recherche du Québec Santé (FRQS, 348782). Trainees were supported by FRQS Doctoral Studentships (to RRJ, DS), Canada Graduate Scholarship – Doctoral Award (to LTT) and Masters Awards (to JT, JEK), Cole Foundation Studentships (to LTT, DS), Cancer Research Society Doctoral Award (to DS), and Fellowship from Arthritis Society Canada and the Richard and Edith Strauss Foundation (to MM). We thank Patricia D’Arcy for mouse management and disease induction, and McGill Comparative Medicine and Animal Resources Centre (CMARC) for mouse husbandry and veterinary services. We thank Sonia Léger-Thériault, Clinical Research Coordinator-Nurse at RI-MUHC, for patient recruitment and blood collection. Flow cytometry was performed at McGill Life Sciences Complex, and we thank facility managers Drs. Camille Stegen and Julien Leconte. We thank Digital Research Alliance of Canada and Calcul Québec for resources for data analyses. Graphical abstract was prepared with BioRender.

## Author Contributions

Data were acquired by JT, RRJ, and LTT, with support from BP and JEK, and with supervision from AN, DL, IC, SMV, DM, MY, and MM. Murine models were established by MY, SMV, and DM. Experiments were designed by AN, JT, RRJ, LTT, DL, IC, SMV, and DM. Data were analyzed by JT, LTT, DS, RRJ, AN. Manuscript was written by AN, JT, RRJ, DS and edited by all co-authors.

---

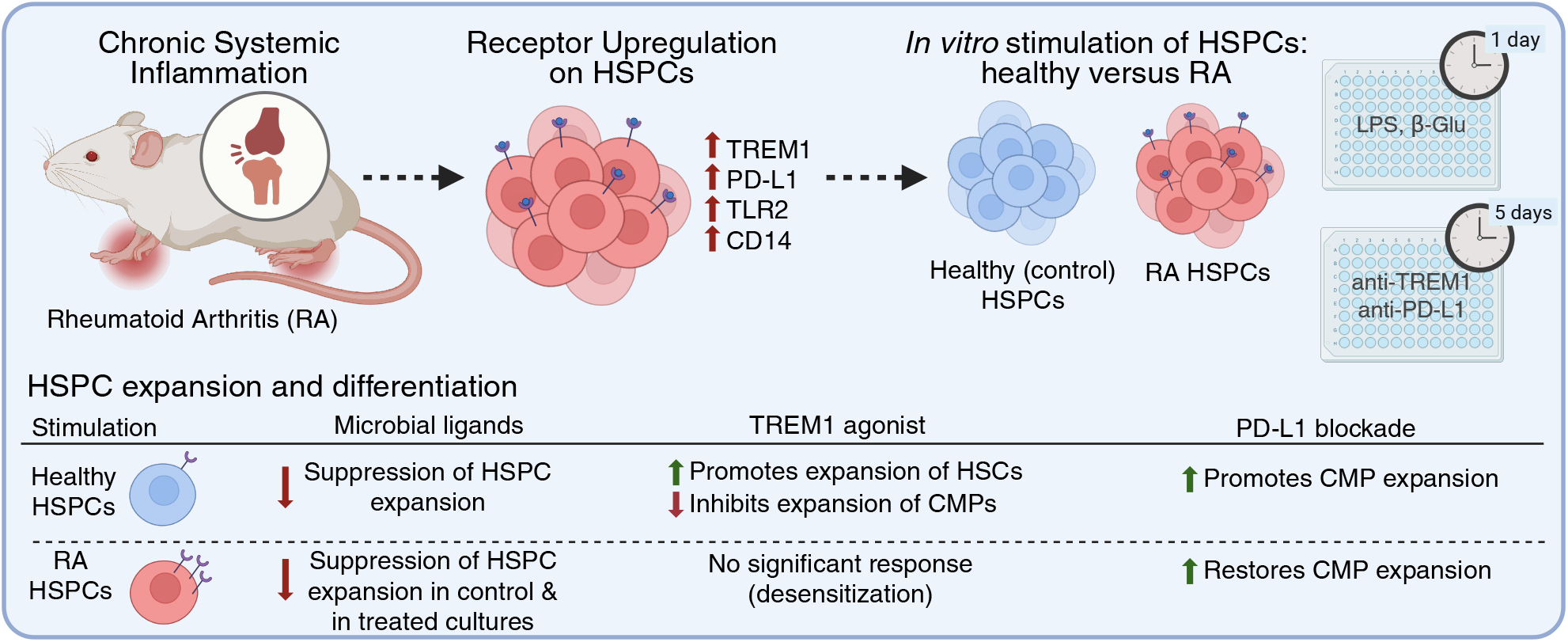

