## Supplemental Materials for "Increased Expression and Altered Functional Activities of Immune Receptors TREM1, PD-L1, and Others on Hematopoietic Progenitor Cells in a Mouse Model of Rheumatoid Arthritis"

### SUPPLEMENTAL MATERIALS AND METHODS

**Mouse Model of Rheumatoid Arthritis:** The SKG mouse model of rheumatoid arthritis was previously described [1], and used extensively in our recent work [2, 3]. The SKG mouse strain carries a homozygous G-to-T substitution at nucleotide 489 in *Zap70* gene on BALB/c genetic background [1]. Rheumatoid arthritis is induced in the mice with a single intraperitoneal injection of zymosan (Millipore-Sigma, 58856-83-2) at 5 mg per mouse at the age of 8-10 weeks [2, 3]. Littermate SKG mice receiving an equal volume of PBS were used as healthy controls. The mice were analyzed at ≥5 weeks after the zymosan treatment when the animals had a clinical score of 8-16, indicating medium to severe arthritis [4]. Mice were maintained under SPF conditions, and matched by age and sex between groups. Mouse experiments were in accordance with guidelines of the Canadian Council on Animal Care and protocols MCGL-7932 and 6029 approved by the McGill Animal Care Committee.

**Blood Mononuclear Cell Isolation:** Human blood was diluted with an equal volume of phosphate-buffered saline pH 7.4 (PBS, Invitrogen) and separated by density gradient centrifugation over Lymphoprep (Stem Cell Technologies), according to manufacturer's protocol. The mononuclear cell layer was collected, washed twice in PBS with 2% (v/v) fetal calf serum (FCS, ThermoFisher Scientific), and counted using a trypan blue exclusion method.

**Flow Cytometry:** Cell suspensions of mouse tissues were prepared in RPMI-1640 (ThermoFisher Scientific) with 2% (v/v) fetal calf serum (FCS, ThermoFisher Scientific), 100 µg/mL streptomycin (Wisent), and 100 U/mL penicillin (Wisent). The samples were stained for cell surface markers in PBS with 2% (v/v) FCS and Super Bright Complete Staining Buffer (SB-4401-75, 1:20, ThermoFisher Scientific) for 20 minutes on ice, using the fluorophore conjugated antibodies provided in Supplemental Tables S1-S3. Viability dyes eFluor 506 or eFluor 780 (ThermoFisher Scientific) were used to discriminate dead cells. Compensation was performed with BD CompBeads (BD Biosciences) or UltraComp eBeads Plus (ThermoFisher Scientific). Data were acquired on BD Fortessa and analyzed with FlowJo software (BD Biosciences).

**RNA sequencing data analyses:** Feature counts from RNA-seq datasets generated by Regan-Komito et. al. [5] were accessed from NCBI public database under GEO accession number GSE126218. Gene expression was quantified based on uniquely mapped reads using featureCounts with default parameters (Subread package v1.5.2) [6]. Genes with counts per million (CPM) greater than 5 in at least 3 samples were retained, with a total of 10,927 genes in LT-HSC/HSC dataset, 10,538 genes in ST-HSC/MPP1-2 dataset, 10,219 genes in MPP/MPP3-4 dataset, and 9,377 genes in GMP dataset; (dual cell names indicate the nomenclature used in the original study and the current work, respectively). TMM normalization and differential expression analysis were carried out using the edgeR Bioconductor package [7]. Genes showing ≥ |1.5|-fold change with a Benjamini–Hochberg adjusted $p$-value ≤ 0.05 were considered significantly differentially expressed.

**Statistics:** Statistical analyses were performed with Prism 9.5.1 or Prism 10.5.0 (GraphPad Inc.), with statistical tests for each dataset indicated in the Figure Legends.

**Table S1. Antibodies for flow cytometry analyses of murine bone marrow.**

| Target | Fluorophore | Company | Clone | Catalogue |
|---|---|---|---|---|
| CD3 | Biotin | Biolegend | 17A2 | 100244 |
| B220 | Biotin | Biolegend | RA3-6B2 | 103203 |
| TER119 | Biotin | Biolegend | TER-119 | 116204 |
| Streptavidin | Brilliant Violet 785 | Biolegend | NA | 405249 |
| cKIT | Brilliant Violet 650 | Biolegend | ACK2 | 135125 |
| cKIT | PE-Cy7 | Biolegend | 2B8 | 105814 |
| SCA1 | Brilliant Ultraviolet 395 | BD Biosciences | D7 | 563990 |
| SCA1 | APC | eBioscience, Thermo Fisher | D7 | 17-5981-83 |
| FLT3 | PE | eBioscience, Thermo Fisher | A2F10 | 12-1351-83 |
| CD34 | Brilliant Violet 421 | Biolegend | SA376A4 | 152208 |
| CD150 | PE-Cy7 | Biolegend | TC15-12F12.2 | 115913 |
| CD48 | PerCP-Cy5.5 | Biolegend | HM48-1 | 103422 |
| CD16/CD32 | Brilliant Ultraviolet 737 | BD Biosciences | 2.4G2 | 612783 |
| CD16/CD32 | FITC | eBioscience, Thermo Fisher | 93 | 11-0161-85 |
| TREM1 | eFlour660 | eBioscience, Thermo Fisher | TR3MBL1 | 50-3541-80 |
| Dectin1 | FITC | eBioscience, Thermo Fisher | 2A11 | MA5-16479 |
| TLR2 | APC | Biolegend | QA16A01 | 153006 |
| CD14 | FITC | Biolegend | Sa14-2 | 123308 |
| CD45 | Brilliant Ultraviolet 395 | BD Biosciences | 30-F11 | 564279 |

**Table S2. Antibodies for flow cytometry analyses of human HSPCs.**

| Target | Fluorophore | Company | Clone | Catalogue |
|---|---|---|---|---|
| CD45 | Brilliant Ultraviolet 395 | BD Biosciences | HI30 | 563792 |
| CD34 | PE-Cy7 | Biolegend | 581 | 343515 |
| CD38 | Brilliant Violet 785 | Biolegend | HIT2 | 303530 |
| TLR2 | FITC | Biolegend | TL2.1 | 309705 |
| TREM1 | APC | Biolegend | TREM-26 | 314909 |
| CD14 | PE | Biolegend | M5E2 | 301806 |

**Table S3. Antibodies for flow cytometry analyses of murine HSPC cultures.**

| Target | Fluorophore | Company | Clone | Catalogue |
|---|---|---|---|---|
| CD3 | Biotin | BioLegend | 17A2 | 100244 |
| B220 | Biotin | BioLegend | RA3-6B2 | 103204 |
| TER119 | Biotin | BioLegend | TER-119 | 116204 |
| Ly6G | Biotin | BioLegend | 1A8 | 127604 |
| Streptavidin | Brilliant Violet 785 | BioLegend | N/A | 405249 |
| CD45 | Spark PLUS Ultraviolet 395 | BioLegend | 30-F11 | 103192 |
| CD117 (cKit) | PE | BioLegend | 2B8 | 105807 |
| Sca-1 | Alexa Fluor 488 | BioLegend | E13-161.7 | 122516 |
| CD34 | PE-Cy7 | BioLegend | SA376A4 | 152218 |
| CD135 (Flt3) | Brilliant Violet 421 | BioLegend | A2F10 | 135315 |
| CD16/32 | Brilliant Ultraviolet 737 | BD Biosciences | 2.4G2 | 612783 |
| CD150 | Brilliant Violet 650 | BioLegend | TC15-12F12.2 | 115932 |
| CD48 | PerCP-Cy5.5 | BioLegend | HM48-1 | 103422 |
| CD71 | APC-Fire750 | BioLegend | R17217 | 113828 |

**Table S4. Demographic characteristics of patients recruited for the study.**

| | |
|---|---|
| Age | 39 y, 51 y, 68 y |
| Sex | 2 F, 1M |
| Comorbidities | |
| Smoking | - |
| Hypertension | 1 |
| Medication | Two patients were unmedicated, and one received methotrexate (15mg/week) & hydroxychloroquine (400mg/day). |

### SUPPLEMENTAL DATA FIGURES

**Figure S1. Expression of TREM1, TLR2, and CD14 on hematopoietic stem and progenitor cells (HSPCs) of human arthritis patients and healthy controls, matched by age and sex.** Peripheral blood was processed to isolate the mononuclear cell fraction and HSPCs were gated as $CD34^{+}$ cells. Statistical analyses used paired *t*-test, due to the technical pairing of samples processed on the same day; * $p$<0.05, ** $p$<0.01; MFI – mean fluorescence intensity.

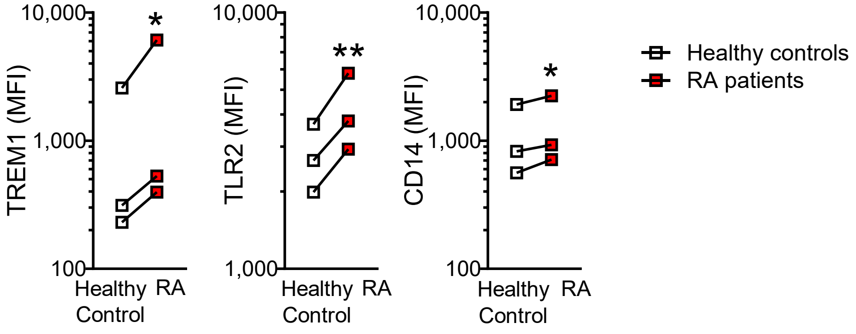

**Figure S2. Select differentially expressed genes encoding immune receptors in hematopoietic stem and progenitor cells of SKG mice with systemic inflammatory disease.** RNA-Seq data generated by Regan-Komito et. al. [5] were accessed from NCBI public database under GEO accession number GSE126218 and re-analyzed for this study. The four populations analyzed in the study are: $Lin^{-}$ $cKit^{+}Sca1^{+}CD34^{-}CD150^{+}CD48^{-}$ (LT-HSC, referred to here as HSC); $Lin^{-}cKit^{+}Sca1^{+}CD34^{+}CD150^{+}$ (ST-HSC, referred to here as MPP1-2); $Lin^{-}cKit^{+}Sca1^{+}CD34^{+}CD150^{-}CD48^{+}$ (MPP, referred to here as MPP3-4), and $Lin^{-}cKit^{+}Sca1^{-}CD34^{+}CD16/32^{hi}$ (GMP). Fold changes indicate the expression of each gene in SKG mice with systemic inflammation relative to its expression in healthy control SKG mice for each cell type. Select genes that encode the receptors studied in our work or closely related receptors are highlighted.

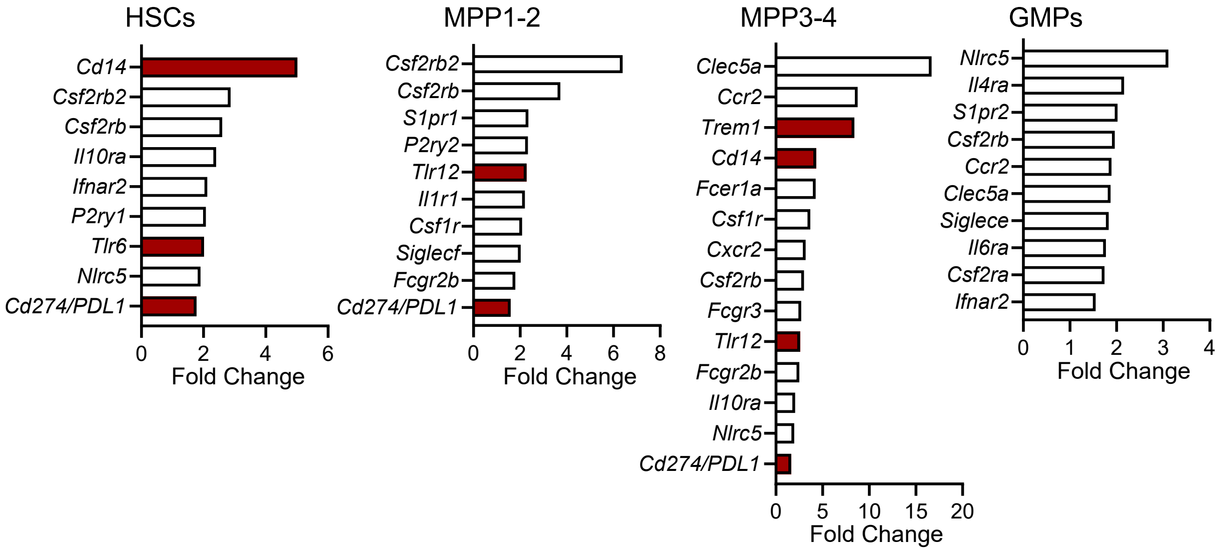

**Figure S3. Gating of murine hematopoietic stem and progenitor cells. (A-B)** Gating on live lineage negative hematopoietic stem and progenitor cells, corresponding to Lin$^{-}$cKit$^{+}$Sca1$^{+}$ (LKS) cells and Lin$^{-}$cKit$^{+}$Sca1$^{-}$ (LK) cells. **(C)** Gating on common myeloid progenitors (CMPs), granulocyte-monocyte progenitors (GMPs), and megakaryocyte-erythroid progenitors (MEPs) within LK cell population, based on CD34 and CD16/32 markers. **(D-E)** Gating on hematopoietic stem cells (HSCs) and multipotent progenitors 1-4 (MPP1-4) within the LKS cell population, based on CD150, CD48, CD34, and CD135/FLT-3 markers. Gating strategies are widely established, and used extensively in our recent work [8, 9].

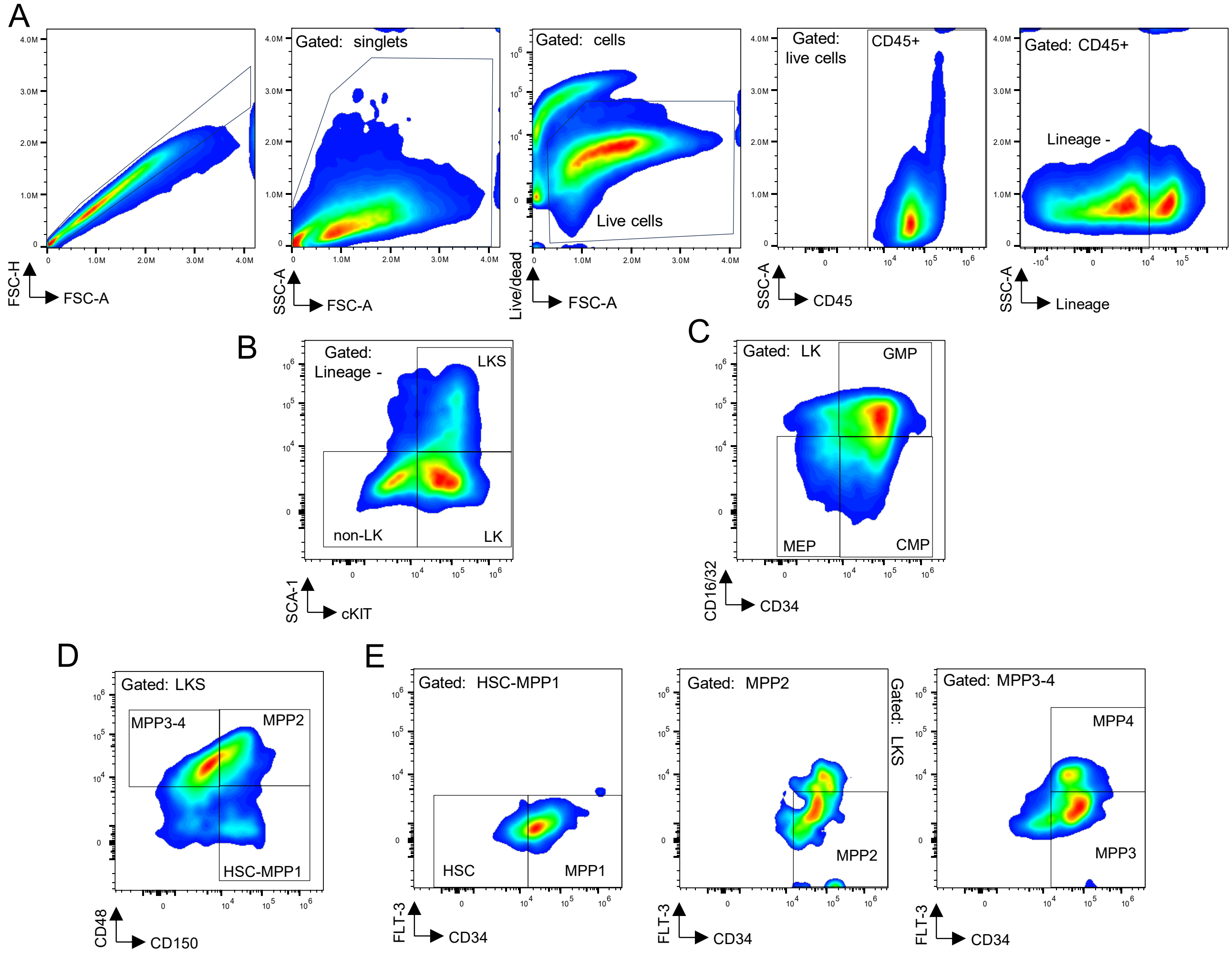

**SUPPLEMENTAL REFERNCES**
